# Mercator: An R Package for Visualization of Distance Matrices

**DOI:** 10.1101/733261

**Authors:** Zachary B. Abrams, Caitlin E. Coombes, Suli Li, Kevin R. Coombes

**Author notes:** These authors contributed equally to this work (Zachary B. Abrams, Caitlin E. Coombes).

## Abstract

**Summary:** Unsupervised data analysis in many scientific disciplines is based on calculating distances between observations and finding ways to visualize those distances. These kinds of unsupervised analyses help researchers uncover patterns in large-scale data sets. However, researchers can select from a vast number of different distance metrics, each designed to highlight different aspects of different data types. There are also numerous visualization methods with their own strengths and weaknesses. To help researchers perform unsupervised analyses, we developed the Mercator R package. Mercator enables users to see important patterns in their data by generating multiple visualizations using different standard algorithms, making it particularly easy to compare and contrast the results arising from different metrics. By allowing users to select the distance metric that best fits their needs, Mercator helps researchers perform unsupervised analyses that use pattern identification through computation and visual inspection.

**Availability and Implementation:** Mercator is freely available at the Comprehensive R Archive Network (https://cran.r-project.org/web/packages/Mercator/index.html)

**Supplementary information:** Supplementary data are available at *Bioinformatics* online.

## 1 Introduction

Data visualization is a key component of scientific analysis. Observing visual patterns in a data set aids researchers in understanding the relationships among their data as they search for greater biological meaning. As more experiments are conducted on increasingly large-scale, multi-dimensional data sets, new visualization methods must be developed to keep pace with the technical demands and research potential of these more complicated data sets. This problem is especially acute in biomedical research with the advent of “omics” technologies that can measure tens of thousands of features across thousands of samples.

Many unsupervised analyses rely on calculating similarities and differences between the observed samples based on a large feature space. Since many biomedical experiments seek to define separations or distinctions between groups, they must measure that difference using some distance metric. Based on the type of data (e.g., binary or continuous), the appropriate distance metric must be chosen to quantify a separation between data objects, based on the best fit for specific data types and experimental criteria. Thus, there is not one distance metric that is seen as superior to all others in all contexts. Here we present Mercator, a new R package for visualizing distance matrices.

## 2 Methods

Mercator is designed to calculate distance matrices and visualize data based on the user’s chosen methods. To begin, Mercator takes the user input in the form of a data matrix of binary or continuous variable data. Next, initial data filtering may be performed using Thresher (Wang, et al., 2018), an R package that performs clustering using a combination of outlier detection, principal components analysis, and von Mises Fisher mixture models. By identifying significant features, Thresher performs feature reduction through removal of uninformative features. It then calculates of the number of groups (K) for downstream use. After that, the user calculates the appropriate distance metric based on data type and biological meaning. Users can use any of the continuous metrics supported by `dist` or similar functions in R. In addition, Mercator implements several distance metrics between binary vectors (Choi, et al., 2010), including Jaccard, Sokal-Michener, Hamming, Russell-Rao, Pearson, and Goodman-Kruskal. Finally, Mercator provides access to four visualization methods, including hierarchical clustering, multi dimensional scaling (MDS) (Cox and Cox, 2001), t-distributed Stochastic Neighbor Embedding (t-SNE) (van der Maaten and Hinton, 2008), and iGraph (Csardi and Nepusz, 2006). Users may easily mix and match distance metrics and visualization techniques to gain a better understanding of patterns in their data; Mercator includes tools to ensure that colors and symbols are as consistent as possible when multiple distance metrics are applied to the same data set.

An example is shown in Figure 1. Cytogenetic data obtained from the Mitelman Database of Chromosomal Aberrations and Gene Fusions in Cancer (Mitelman, et al., 1991) for Chronic Lymphocytic Leukemia (CLL) patients was processed using CytoGPS (Abrams, et al., 2019) and converted into a binary matrix. After data filtering with Thresher, a Jaccard distance matrix was calculated and visualized with MDS, t-SNE, and iGraph. The t-SNE plot highlights both clearly defined clusters and closely located groups. MDS plots expand this understanding by separation by the most important, distinguishing component between features; higher dimensional MDS plots are required to elucidate the entire structure. After down-sampling the data set to underemphasize the strongest signals while preserving the weaker ones, the iGraph visualization provides a better understanding of interactions between the cytogenetics of CLL.

**Figure 1:**
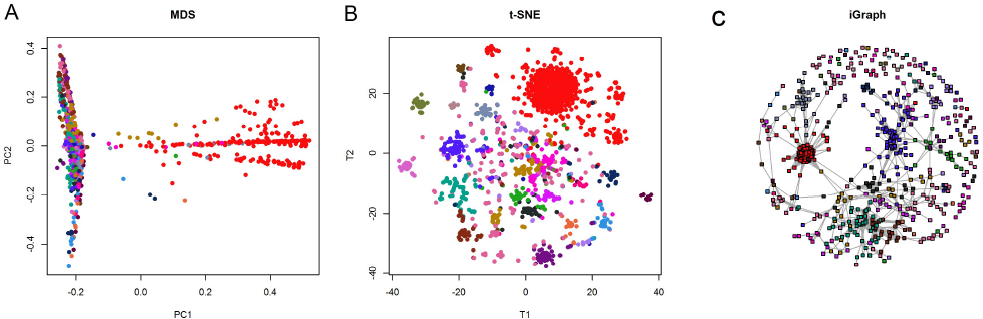
Visualizations of CLL cytogenetic data. (A) The first coordinate of the multidimensional scaling plot shows the dominant signal (trisomy 12) in samples colored red. (B) The t-SNE plot shows not only that signal, but clearly reveals other clusters based on patterns of cytogenetic abnormalities. (C) The down-sampled iGraph representation gives a better idea of the connections between clusters.

## 3 Conclusion

Mercator is in R package that visualizes large-scale, multi-dimensional data using 10 distance methods and 5 visualization techniques. The package facilitates easy visualization and comparison to aid greater understanding the underlying patterns present in their data.

## Acknowledgements

We would like to acknowledge the support from the National Library of Medicine and from the Summer Internship Program at the Ohio State University Department of Biomedical Informatics.

## Funding

This work was supported by the National Library of Medicine (NLM) [grant number T15 LM011270], the National Cancer Institute (NCI) [grant number R03 CA235101], and by Pelotonia Intramural Research Funds from the James Cancer Center, Columbus, OH.

## Notes

https://cran.r-project.org/package=Mercator

